# Redundancy in citrate and *cis*-aconitate transport in *Pseudomonas aeruginosa*

**DOI:** 10.1101/2022.07.01.498397

**Authors:** Simon A.M. Underhill, Matthew T. Cabeen

## Abstract

Tricarboxylates such as citrate are the preferred carbon sources for *Pseudomonas aeruginosa*, an opportunistic pathogen that causes chronic human infections. However, the membrane transport process for the TCA cycle intermediates citrate and *cis*-aconitate is poorly characterized. Transport is thought to be controlled by the TctDE two-component system, which mediates transcription of the putative major transporter OpdH. Loss of *tctDE* has been associated with sensitization to aminoglycosides, possibly linking tricarboxylate transport to enhanced antimicrobial resistance. In this work, we search for previously unidentified transporters of citrate and *cis*-aconitate using both protein homology and RNA sequencing approaches. We uncover new transporters and show that OpdH is not the major citrate porin; instead, citrate transport primarily relies on the tripartite TctCBA system, which is encoded in the *opdH* operon. Deletion of *tctA* causes a growth lag on citrate and loss of growth on *cis*-aconitate. Combinatorial deletion of newly discovered transporters can fully block citrate utilization. We then characterize transcriptional control of the *opdH* operon in *tctDE* mutants and show that loss of *tctD* blocks citrate utilization due to its inability to express *opdH-tctCBA*. However, *tctE* and *tctDE* mutants evolve heritable adaptations that restore growth on citrate as the sole carbon source.

**Author Summary:** *Pseudomonas aeruginosa* is a bacterium that infects hospitalized patients and is often highly resistant to antibiotic treatment. It preferentially uses small organic acids called tricarboxylates rather than sugars as a source of carbon for growth. The transport of many of these molecules from outside the cell to the interior occurs through unknown channels. In some cases, there may be links between antibiotic uptake and the transport of metabolic molecules, making cross-membrane transport medically important. In this work we examined how the tricarboxylates citrate and *cis*-aconitate are transported in *P. aeruginosa*. We then sought to understand how production of proteins that permit citrate and *cis*-aconitate transport is regulated by a signaling system called TctDE. We identified new transporters for these molecules, clarified the function of a known transport system, and directly tied transporter expression to the presence of an intact TctDE system.

## Introduction

The Gram-negative bacterium *Pseudomonas aeruginosa* thrives on many carbon sources, but unlike *Escherichia coli* and many other model bacterial species shows a preference for TCA cycle intermediates such as succinate or citrate over glucose and other saccharides [1]. The import, modification, and metabolism of glucose in *P. aeruginosa* has been extensively studied as a case of a bacterium that does not possess the textbook Embden-Meyerhof-Parnas (EMP) glycolytic pathway [2]. This organism also does not possess sugar phosphotransferase systems (PTS) for many of the carbon sources it uses, but rather a series of OprD family outer membrane porins that are typically induced by their respective substrates [3]. The use of specific porins by *P. aeruginosa* has been linked to lower membrane permeability and, along with the presence of efflux pumps, bacterial tolerance to antibiotic treatment [4]. Because *P. aeruginosa* is responsible for many hospital-acquired infections [5] and chronically infects the lungs of cystic fibrosis patients [6, 7], it is important to characterize membrane transport as a step toward learning how transport impacts antibiotic tolerance.

Despite being a TCA cycle intermediate and hence a preferred carbon source for *P. aeruginosa* [8–10], citrate transport is not well understood in this organism. A transposon-insertion mutant of the *opdH* porin-encoding gene showed a growth defect on *cis*-aconitate [3], a molecular neighbor to citrate in the TCA cycle. The *opdH* gene was additionally induced on citrate but not glucose, but a mutant showed no growth defect on citrate [11]. OpdH is a predicted porin and showed channel-like activity in a planar bilayer apparatus, but transport of citrate or *cis*-aconitate could not be demonstrated in these experiments [11]. The *opdH* gene is in an operon with the *tctCBA* genes, which encode a predicted tripartite tricarboxylate transport system. This system has not previously been tested for transport of citrate in *P. aeruginosa* [11], though the homologous TctC subunit is known to bind citrate, isocitrate, and *cis*-aconitate in *Salmonella enterica* [12]. The TctA and TctB proteins in *Salmonella typhimurium* are predicted transmembrane proteins that are putatively involved in the cross-membrane transport of the substrate bound by TctC [13]. In *P. aeruginosa,* the *opdH*-*tctCBA* operon is divergently transcribed from the *tctDE* operon, which encodes a two-component system (TCS) that is thought to repress *opdH-tctCBA* expression via binding of the TctD response regulator to the promoter region [11, 14]. A *tctDE* mutant reportedly failed to grow on citric acid as a sole carbon source [15], which raises the question of how the citrate-induced *opdH-tctCBA* operon behaves in citrate in the absence of its repressor. If TctDE is a simple TCS that senses TCA cycle intermediates and responds by de-repressing *opdH-tctCBA*, loss of the TctD repressor would result in constitutive expression of those genes rather than loss of growth in citrate. The *opdH-tctCBA* operon also appears to encode another protein, PA14_54580, which has not been previously examined. PA14_54580 is annotated as a conserved hypothetical protein of unknown function, though one report [11] labeled it as a monooxygenase.

In *S. typhimurium,* a citrate transporter named CitA was characterized and could facilitate transport of citrate in *E. coli* cells [16], which do not normally grow on citrate, as they lack a functional transporter [17–19]. A previously uncharacterized homolog of the *S. typhimurium citA* gene exists in *P. aeruginosa* in both PAO1 and PA14 (*PA5476* in PAO1; *PA14_72280* in PA14). While it was suggested that *opdH* may be required for citrate transport [15], the presence of a *citA* gene that is highly similar to a predicted major facilitator superfamily (MFS) transporter of *S. enterica* (59.9% identity, 74% similarity by the EMBOSS Needle alignment tool [20]) and *citA* of *S. typhimurium* (59% identity, 72.5% similarity) suggests that citrate transport in *Pseudomonas* is more redundant than currently believed. However, the predicted *citA* transporter gene had not yet been assessed for expression or function.

In this work, we examine redundancy in and control of citrate and *cis*-aconitate transport in *P. aeruginosa* by using a series of genetic deletions coupled with growth screens and by complementing nonfunctional citrate and *cis*-aconitate transport in *E. coli* using transporters from *Pseudomonas*. We clarify the role of OpdH, previously thought to be the major porin for citrate [15], and identify several new transport proteins using transcriptomic analysis. We then use deletions of the *tctDE* TCS coupled with RT-qPCR to probe the impact of TctDE on expression of citrate transporter-encoding genes in the presence and absence of citrate.

## Results

### *opdH* and/or *citA* deletions do not affect growth on citrate but modestly impact *cis*-aconitate growth

Having identified *citA* as a gene encoding a putative citrate transporter, we constructed strains to test whether deletion of *citA* would affect the ability of *P. aeruginosa* to grow on citrate. We obtained growth curves by measuring the OD_600_ of *P. aeruginosa* PA14 derivative cultures in M9 minimal medium supplemented with citrate as the sole carbon source. The Δ*citA* and Δ*opdH* strains showed no obvious growth defects relative to the wild type (Fig. 1A), in contrast to previous results using an *opdH* gentamicin cassette insertion mutant in strain PAO1 that resulted in a modest citrate growth defect and a severe aconitate growth defect [11]. A Δ*citA* Δ*opdH* double mutant showed a marginally longer lag and a slightly slower initial growth rate that were barely noticeable (Fig. 1A). On *cis*-aconitate as the sole carbon source, we again observed no growth defects for the single Δ*citA* and Δ*opdH* strains (Fig. 1B). This result also stands in contrast to previous reports of severe *cis*-aconitate growth defects in a transposon insertion or Gm^R^ insertion *opdH* mutant of strain PAO1 [3, 11]. However, the Δ*citA* Δ*opdH* double mutant displayed a substantial lag in growth on *cis*-aconitate (∼ 500 minutes vs. ∼ 200 minutes for the parent and single deletions; Fig. 1B), suggesting that these transporters are more important for *cis*-aconitate transport than for citrate transport and are not the sole transporters of either molecule.

**Figure 1:**
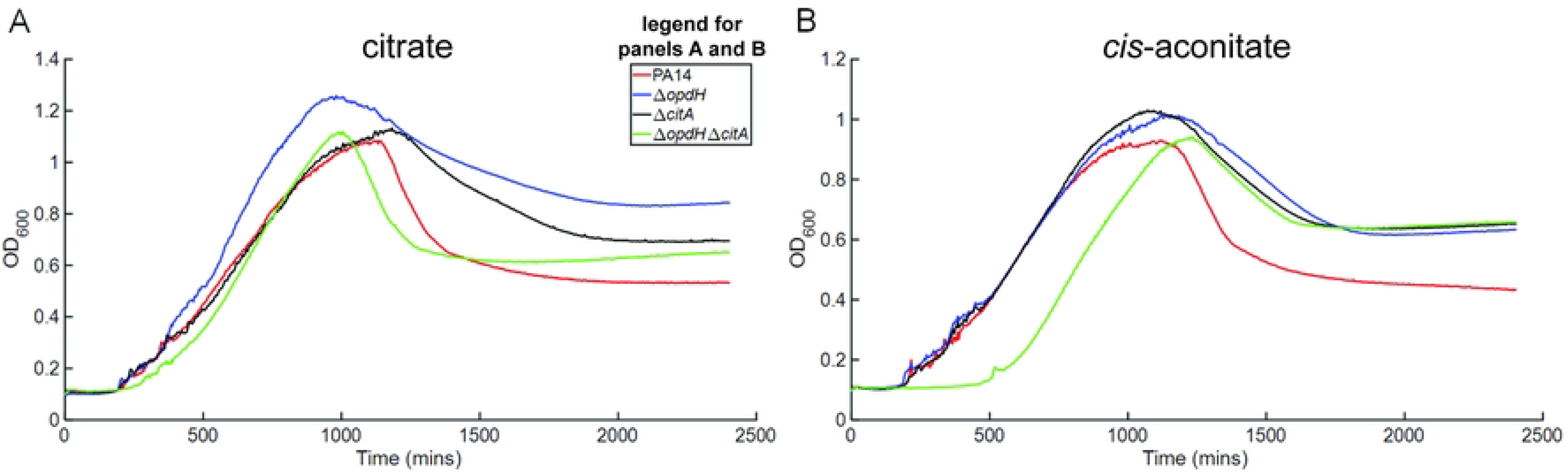
Effects of *citA* and *opdH* on citrate and *cis*-aconitate transport. Growth curves showing OD_600_ vs. time for PA14 (red curves), Δ*opdH* (blue curves), Δ*citA* (black curves), or Δ*opdH* Δ*citA* (green curves) in M9 supplemented with (A) 7.5 mM citrate or (B) 7.5 mM *cis*-aconitate as the sole carbon source. Data are representative curves of at least 3 independent experiments.

### CitA, but not OpdH, enables citrate and *cis*-aconitate transport in *E. coli*

*E. coli* MG1655 is a K-12 derivative that, like other K-12 derivatives, does not transport citrate and therefore does not grow on citrate as a sole carbon source. We took advantage of this property to test whether expression of *citA* or *opdH* under an IPTG-inducible promoter would be sufficient to enable transport of citrate. As expected, MG1655 grew in M9 minimal medium supplemented with glucose but did not grow in M9 with citrate as the sole carbon source (Fig. 2A). As a positive control, an MG1655 strain constitutively expressing *citT*, which encodes a citrate transporter, from a plasmid (gift of Profs. Zach Blount and Richard Lenski [21]) achieved substantial growth in citrate (Fig. 2A). Next, we individually expressed *citA* or *opdH* from an IPTG-inducible pTrc99A plasmid in MG1655. Neither plasmid permitted growth without IPTG (Fig. 2A). When induced, *citA*, but not *opdH*, enabled modest growth on citrate (Fig 2A). We also performed the same complementation test in M9 containing *cis-*aconitate, which like citrate is also not utilized by the MG1655 parent (Fig 2B). Interestingly, expression of *citA* enabled substantial growth of *E. coli* on *cis*-aconitate, whereas expression of either *opdH* or *citT* did not allow growth (Fig. 2B). These results imply that CitA (unlike CitT) can transport both citrate and aconitate but prefers aconitate.

**Figure 2:**
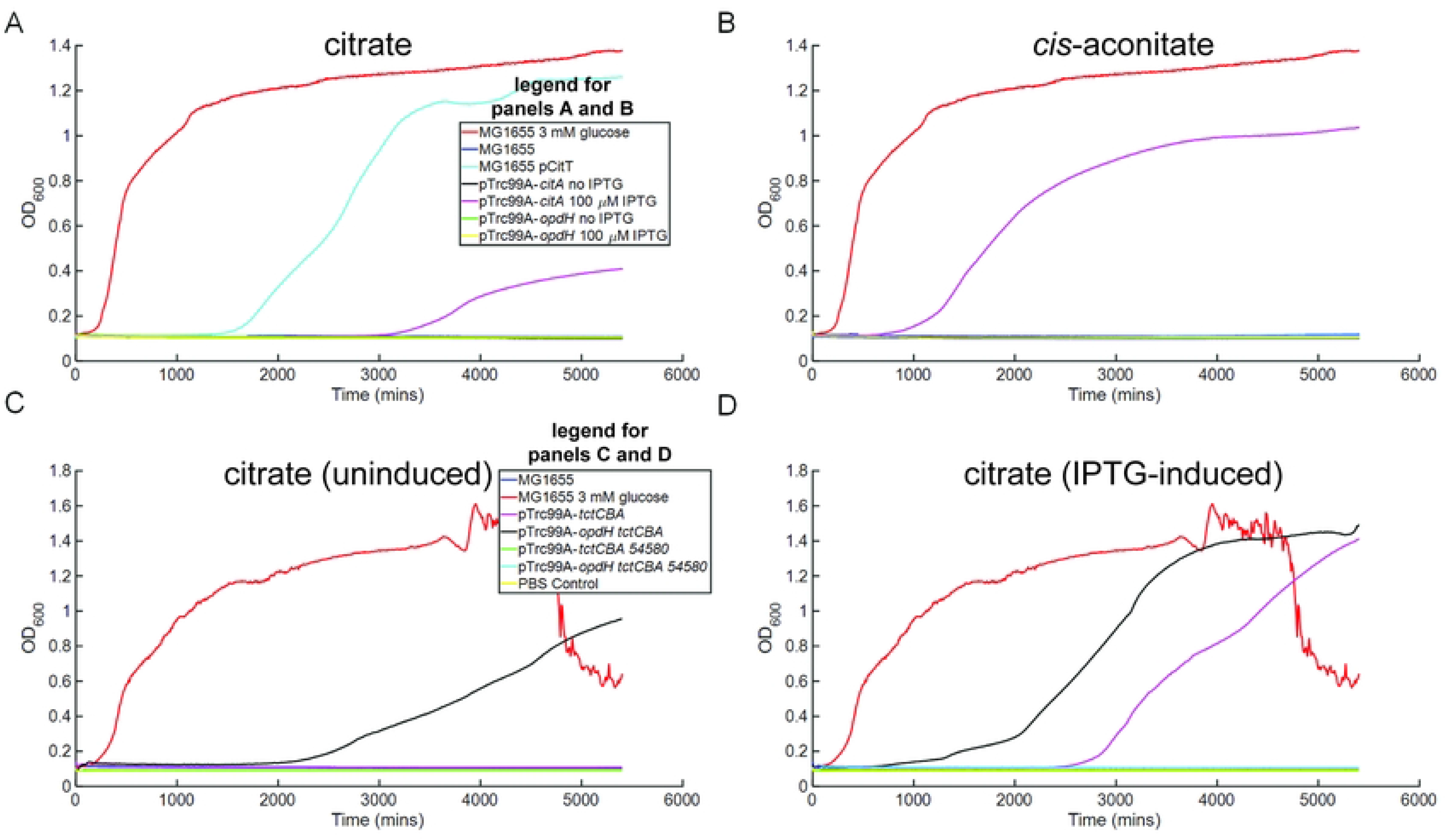
Expression of *citA*, but not *opdH*, facilitates growth of *E. coli* on citrate and *cis*-aconitate. Growth curves showing OD_600_ vs. time for *E. coli* MG1655 bearing different transporter-expressing plasmids growing in M9 supplemented with 7.5 mM citrate or 7.5 mM *cis* aconitate as the sole carbon source. In all panels, MG1655 growing in 3 mM glucose as a positive control is shown by the red curves. (A), (B) Growth in citrate (A) or *cis*-aconitate (B) of MG1655 (blue curves); MG1655 with the CitT positive control transporter expressed (cyan curves); MG1655 carrying pTrc99A with the *citA* CDS with (magenta curves; 100 µM) and without (black curves) IPTG inducing expression of *citA*; and MG1655 carrying pTrc99A with the *opdH* CDS with (yellow curves; 100 µM) and without (green curves) IPTG inducing expression of *opdH*. (C), (D) Growth in M9 supplemented with citrate without (C) or with (D) induction (100 µM) of IPTG in strains bearing fragments of the *opdH* operon. Blue curves, MG1655 wild type; magenta curves, MG1655 expressing the *tctCBA* segment; black curves, MG1655 expressing the *opdH-tctCBA* segment; green curves, MG1655 expressing the *tctCBA-PA14_54580* segment; cyan curves, MG1655 expressing the full *opdH* operon; yellow curves, control in which 2 µL of the PBS used to wash cells was inoculated in the M9 to show sterility. Data are representative curves of at least 3 independent experiments.

Given that previous insertional mutants of *opdH* in PAO1 [3, 11] resulted in citrate and aconitate growth defects, and because OpdH alone did not enable citrate or aconitate utilization (Fig. 2A-B), we considered the possibility that the downstream *tctCBA* genes (which might have been subject to polar effects by insertional mutation) have a role in tricarboxylate transport. Thus, we placed different sections of the *opdH* operon on pTrc99A for expression in *E. coli* and growth analyses. Strikingly, even in the absence of IPTG induction, the *opdH-tctCBA* construct – encoding the TctCBA tripartite transporter and OpdH porin – grew slowly, whereas none of the other tested fragments (*tctCBA, tctCBA-PA14_54580,* and *opdH-tctCBA-54580*) grew (Fig. 2C). When induced, the *opdH-tctCBA* construct grew more quickly than when uninduced, and the construct encoding only the putative TctCBA transporter also grew, though less well than the construct that also included *opdH* (Fig 2D). Interestingly, the constructs bearing *PA14_54580*, a gene of unknown function, failed to permit growth in citrate irrespective of induction despite the presence of *tctCBA* and *opdH* (Fig. 2C-D). Evidently, expression of *PA14_54580* prevents growth of *E. coli* by an unknown mechanism when co-expressed with these transporters.

### Identification of new citrate and aconitate transporters in *P. aeruginosa*

Our findings that *citA* and/or *opdH* deletion do not abolish growth on citrate or *cis-*aconitate (Fig. 1) indicate that there must be at least some redundancy in citrate and *cis*-aconitate transport in *P. aeruginosa*. Furthermore, the complementation tests with *E. coli* (Fig. 2) suggest that OpdH, previously thought to be the major transporter of citrate and *cis*-aconitate in *P. aeruginosa*, does not act on its own but likely requires a partner to transport these solutes. To identify previously unknown citrate transporters, we performed transcriptomics to compare PA14 samples grown in either synthetic cystic fibrosis medium 2 (SCFM2) [22] or SCFM2 + 7.5 mM citrate, reasoning that transporters are often induced by their substrate. In Table 1, we show some of the most citrate-upregulated genes that were also marked as transporters; in principle, some of these may transport citrate and/or *cis*-aconitate in PA14. Indeed, this list of genes includes *opdH* and *tctA*.

**Table 1:** selected genes most upregulated on SCFM2 + 7.5 mM citrate compared to SCFM2 without citrate that were also annotated as a type of transporter.

| Locus tag | Gene name | Annotation | False discovery rate | Log<sub>2</sub> fold change |
| --- | --- | --- | --- | --- |
| *PA14_18320* | N/A | EamA family transporter | $1.63 \times 10^{-16}$ | 5.73 |
| *PA14_54520* | *opdH* | OprD family porin | $2.34 \times 10^{-17}$ | 4.96 |
| *PA14_72170* | N/A | CitMHS family transporter | $4.83 \times 10^{-25}$ | 4.49 |
| *PA14_54570* | *tctA* | tripartite tricarboxylate transporter permease | $1.49 \times 10^{-37}$ | 3.63 |
| *PA14_47560* | N/A | MFS transporter | $6.01 \times 10^{-13}$ | 2.27 |
| *PA14_17610* | N/A | spermidine/putrescine ABC transporter substrate-binding protein | $1.08 \times 10^{-12}$ | 2.55 |
| *PA14_17620* | N/A | ABC transporter permease | $2.84 \times 10^{-14}$ | 3.16 |
| *PA14_17630* | N/A | ABC transporter permease | $2.42 \times 10^{-14}$ | 2.86 |
| *PA14_17640* | N/A | ABC transporter ATP-binding protein | $1.78 \times 10^{-16}$ | 2.52 |
| *PA14_33910* | N/A | ABC transporter substrate-binding protein | $1.46 \times 10^{-17}$ | 2.56 |
| *PA14_21150* | N/A | ABC transporter permease | $2.27 \times 10^{-9}$ | 1.84 |
| *PA14_72960* | N/A | MFS transporter | $4.59 \times 10^{-11}$ | 1.75 |
| *PA14_19160* | *rarD* | EamA family transporter RarD | $8.62 \times 10^{-12}$ | 1.60 |

Guided by our transcriptomic data, we deleted selected suspected transporters to assess their function in citrate or aconitate utilization. Deletions of the *tctA* gene (*PA14_54570*), the two identified genes encoding EamA-family proteins (*PA14_18320* and *rarD*), and *PA14_72170* were all assayed for growth defects on citrate in liquid M9 culture. We found that deletion of *18320, 72170* or *rarD* alone had no significant impact on growth in citrate (Fig. 3A) or *cis*-aconitate (Fig. 3B). In contrast, the Δ*tctA* strain showed a substantially longer lag time in citrate (Fig. 3A) and completely failed to grow on *cis*-aconitate (Fig. 3B). All of these mutants grew similarly to PA14 on M9 with 3 mM glucose (Supp. Fig. S1), implying that the growth changes are specific to tricarboxylate carbon sources.

**Figure 3:**
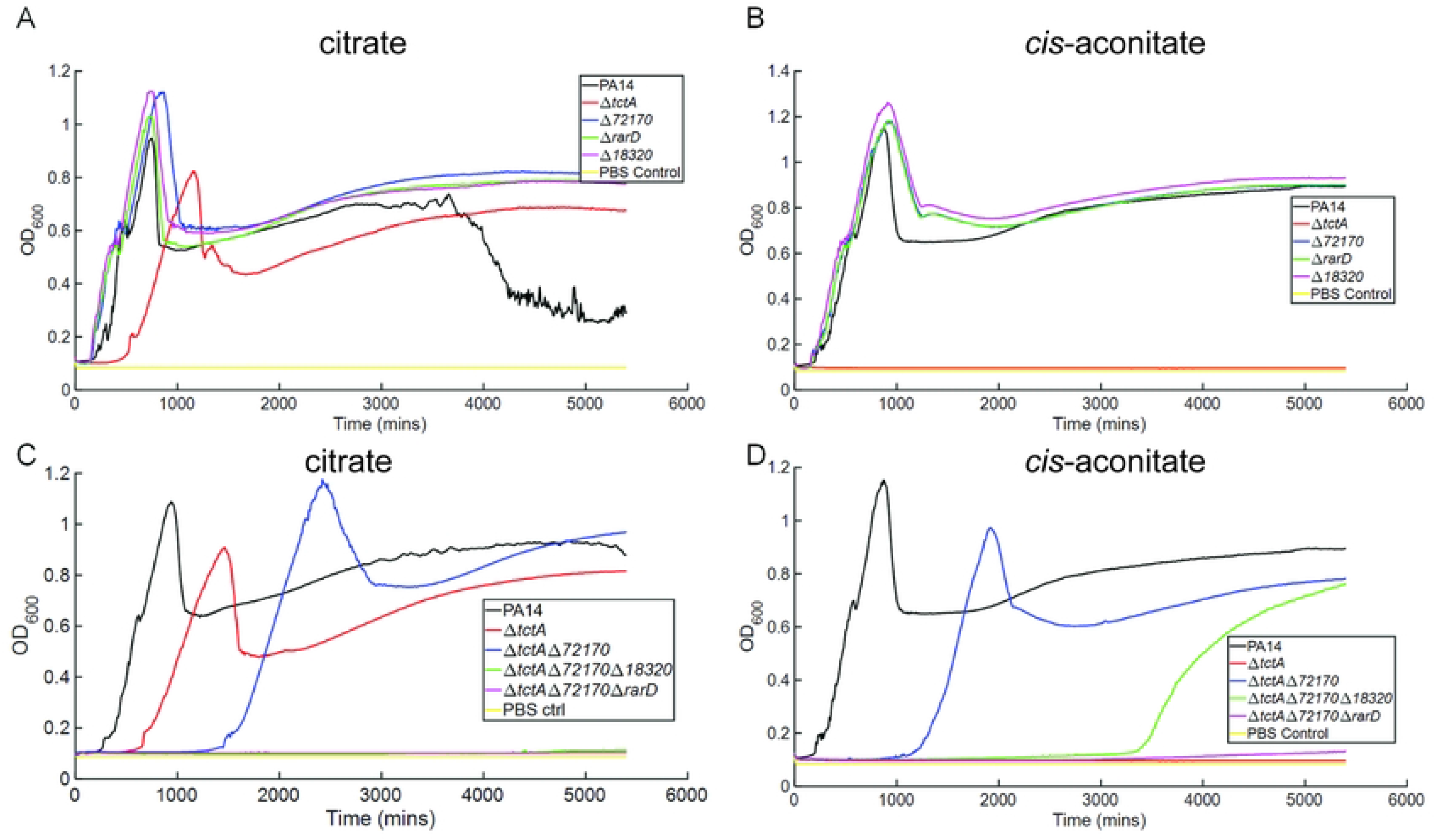
Effect of deletion of newly identified transporters on citrate and *cis*-aconitate growth. Growth curves showing OD_600_ vs. time for mutants of putative new citrate transport proteins. Growth in M9 supplemented with 7.5 mM citrate (A) or *cis*-aconitate (B) as the sole carbon source is shown for PA14 wild type (black curves); Δ*tctA* (red curves); Δ*PA14_72170* (blue curves); Δ*rarD* (green curves); or Δ*PA14_18320* (magenta curves). Yellow curves, control in which 2 µL of the PBS used to wash cells was inoculated in the M9 to show sterility. Growth in 7.5 mM citrate (C) or *cis*-aconitate (D) as the sole carbon source is shown for multiple mutations of these transporters. Black curves, PA14; red curves, Δ*tctA*; blue curves, Δ*tctA* Δ*PA14_72170*; green curves, Δ*tctA* Δ*PA14_72170* Δ*PA14_18320*; magenta curves, Δ*tctA* Δ*PA14_72170* Δ*rarD*. Yellow curves, control in which 2 µL of the PBS used to wash cells was inoculated in the M9 to show sterility. Data are representative curves of at least 3 independent experiments.

We then examined combinatorial deletions of *tctA* with the other identified transporters to learn whether we could build a strain that completely failed to grow on citrate. Deletion of *72170* with *tctA* resulted in a greater lag (Fig. 3C), and further deletion of either *18320* or *rarD* completely abolished citrate growth (Fig. 3C). The similar impact of these deletions at least suggests that 18320 and RarD may act together, such that loss of either has the same effect on citrate transport.

The situation appeared more complicated on *cis*-aconitate. While a Δ*tctA* single mutant did not grow, the Δ*tctA* Δ*72170* deletion did grow, but with a substantial lag of about 1,100 minutes relative to the wild type (Fig. 3D). Further deletion of *18320* increased the lag to roughly 3,300 minutes but did not abolish growth, whereas the Δ*tctA* Δ*72170* Δ*rarD* triple mutant grew very poorly on *cis*-aconitate, with a barely detectable increase in OD_600_ at roughly 4,000 minutes (Fig. 3D). The *18320* and *rarD* deletions yielded similar results in a Δ*citA* Δ*opdH* background (Supp. Fig. S2), agreeing with our conclusion from Fig. 2 that CitA and OpdH are not critical to transport.

To test whether any of the newly identified transporters was sufficient for citrate transport, we ectopically expressed each of the same four genes of interest from pTrc99A in *E. coli* MG1655. We found that either without (Fig. 4A) or with IPTG (Fig. 4B), only *72170* supported *E. coli* growth on M9 with citrate. None of the proteins enabled growth in *cis*-aconitate (Fig. 4A-B). This result suggests that the 72170 protein is a standalone citrate (but not aconitate) transporter, in accord with the reduced growth of *P. aeruginosa* on citrate when *72170* was additionally deleted in a Δ*tctA* background (Fig. 3C). Moreover, the failure of either *18320* or *rarD* to support *E. coli* growth in tricarboxylate (Fig. 4) is consistent with a scenario in which their protein products work together, as mentioned above (Fig. 3C).

**Figure 4:**
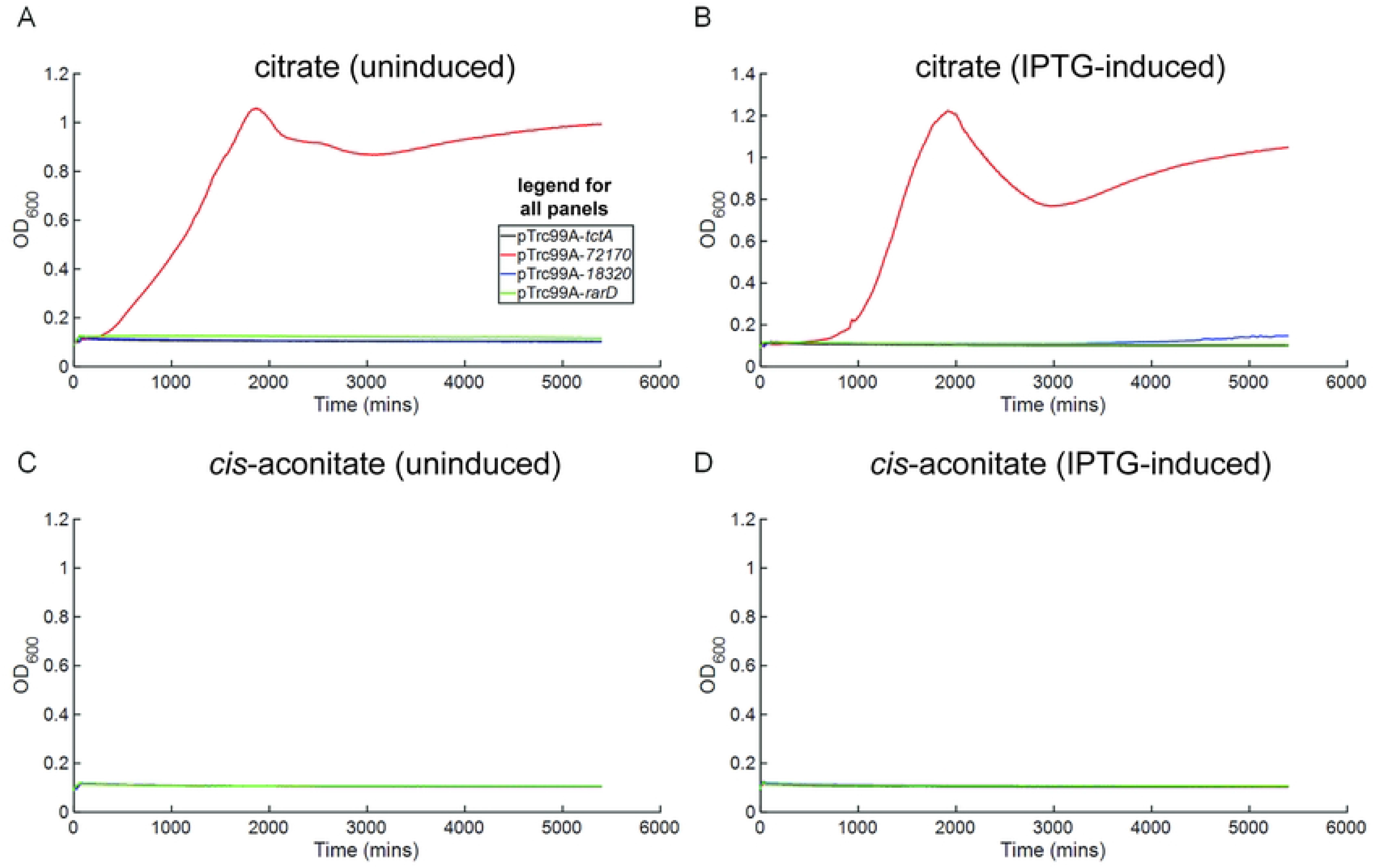
PA14_72170, but not other identified transporters, can facilitate *E. coli* growth on citrate but not *cis*-aconitate. Growth curves showing OD_600_ vs. time for *E. coli* MG1655 carrying pTrc99A constructs encoding putative new citrate transporters. These were tested in M9 supplemented with 7.5 mM citrate as the sole carbon source without (A) or with 100 µM IPTG induction (B); or in 7.5 mM *cis*-aconitate as the sole carbon source without (C) or with 100 µM IPTG induction (D). Black curves, pTrc99A expressing *tctA*; red curves, pTrc99A expressing *PA14_72170*; blue curves, pTrc99A expressing *PA14_18320*; green curves, pTrc99A expressing *rarD*. Data are representative curves of at least 3 independent experiments.

### Deletion of *tctDE* causes a lag in citrate growth followed by heritable adaptation

A previous study found that markerless deletion of the *tctDE* TCS from the genome of *P. aeruginosa* resulted in a growth defect on M63 medium using arginine and citric acid as combined carbon sources [15]. TctD is thought to repress expression of the *opdH* operon in the absence of tricarboxylates [11]; repression is presumably relieved by the TctE kinase when it binds a tricarboxylate substrate extracellularly. We deleted *tctD* and *tctE* independently and in combination to probe how growth in citrate and *cis-*aconitate is affected by each component. Using the previous model as guidance, we expected to lose repression of *opdH* when *tctD* was deleted, resulting in citrate growth. All three *tctD/E* mutants grew normally on glucose (Supp. Fig. S3A) and similarly on a mixture of citrate and glucose (Supp. Fig. S3B), while PA14 grew to a higher OD_600_ on the mixture, consistent with failure of the *tct* mutants to use the citrate. However, we observed that Δ*tctD* mutants were unable to grow in citrate. Surprisingly, the Δ*tctE* and Δ*tctDE* mutants both grew but after a long lag (Fig. 5A; approximately 1,500 minutes for Δ*tctE* and 2,000 minutes for Δ*tctDE*). A control in which 2 µL of the PBS used to wash the cells was inoculated into a well of M9-citrate is shown (Fig. 5A, yellow line) to demonstrate that this late growth was not due to contamination.

**Figure 5:**
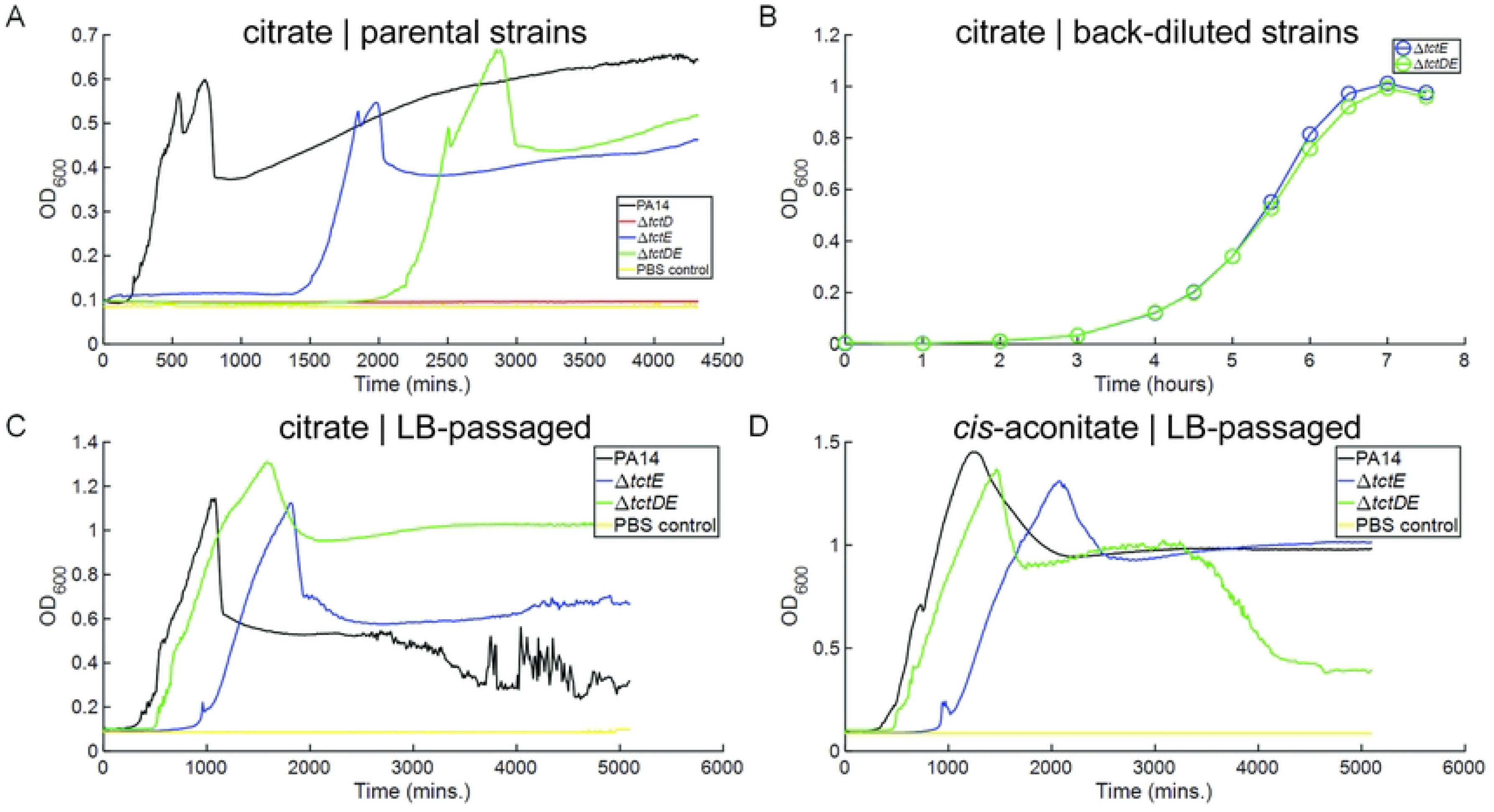
Adaptation of *tctDE* mutants to growth on citrate. Growth curves showing OD_600_ vs. time for *P. aeruginosa* mutants of the TctDE two-component system. (A) Growth data for PA14 (black curve), Δ*tctD* (red curve), Δ*tctE* (blue curve) and Δ*tctDE* (green curve) on M9 supplemented with 7.5 mM citrate as the sole carbon source. (B) The Δ*tctE* (black curve) and Δ*tctDE* (blue curve) strains that grew in panel (A) were back-diluted from M9-citrate culture into M9 with 7.5 mM citrate as the sole carbon source and the OD_600_ vs. time plotted. (C), (D) Growth data for PA14 (black curves), Δ*tctE* (blue curves), and Δ*tctDE* (green curves) passaged on citrate and subsequently streaked on LB plates, grown in liquid LB, and then washed and diluted into M9 supplemented with (C) 7.5 mM citrate or (D) 7.5 mM *cis*-aconitate as the sole carbon source. In panels A, C, and E, the yellow curve represents a control in which 2 µL of the PBS used to wash cells was inoculated in the M9 to show sterility. Data are representative curves of at least 3 independent experiments.

To learn if the growth after the lag was due to acquisition of a heritable adaptation, we grew Δ*tctE* and Δ*tctDE* in shaking flasks in M9-citrate until they were turbid. We then washed these cells twice in PBS and back-diluted them into shaking flasks containing M9-citrate and monitored their growth. Both strains resumed exponential growth after just 3-4 hours (Fig. 5B), a vast reduction compared to their initial lag times and more similar to wild-type cells (Fig. 5A). We then tested whether this adaptation was physiological or genetic by growing three replicates of each background in M9-citrate until they grew, then passaging them on LB agar plates to obtain single colonies and subsequently culturing them in liquid LB. These cells were frozen as stocks for later use, washed twice in PBS, and diluted into M9-citrate and tracked for growth. We present (Fig. 5C) only one replicate as a representative sample. It was immediately evident that the adapted strains grew more quickly in citrate (all start growing after less than 1,000 minutes) than did the parental strains (Fig. 5C). The adapted Δ*tctDE* strain showed the greatest degree of growth acceleration, with a lag of about 500 minutes compared to a parental lag of 2,000 minutes; Δ*tctE* only accelerated from about 1,500 minutes to roughly a 1,000 minute lag time. Because *cis*-aconitate transport appears to overlap significantly with citrate transport, we also tried growing these strains in M9-*cis*-aconitate to see if they had gained a faster ability to grow in *cis*-aconitate. The growth curves are almost identical to those for M9-citrate in lag time (Fig. 5D), showing significant acceleration relative to the parental strains in *cis*-aconitate (Supp. Fig. S4; parental lag times of ∼1000 minutes for Δ*tctE* and ∼2000 minutes for Δ*tctDE*). Together, these data suggest that passaging the Δ*tctE* and Δ*tctDE* strains on citrate as the sole carbon source generates a heritable adaptation to growth on both citrate and aconitate.

### Adapted *tctE* and *tctDE* mutants growing on citrate regain *opdH* expression that is not mimicked by a glucose/citrate mixture

The unexpected heritable adaptation of the *tct* mutants to growth in citrate and aconitate led us to ask how *tctDE* might regulate the expression of the citrate transporters we were studying. To answer this question, we examined transcript levels of the transporter-encoding genes. We grew PA14 and each of our *tct* mutants in M9 supplemented with glucose, citrate, or a mixture of both. We then extracted RNA at mid-exponential phase and performed RT-qPCR. The mixture of glucose and citrate was important for probing expression in the Δ*tctD* mutant, which does not grow on citrate alone. We used the parental (i.e., unadapted) Δ*tctE* and Δ*tctDE* strains for this analysis (note that these strains do not grow on citrate alone until after the long lags described above, implying that adaptation may have taken place).

The first genes we examined by qPCR were *tctD* and *tctE* themselves. This analysis showed that expression of *tctD* and *tctE* was not affected by the carbon source and confirmed that *tctD* and *tctE* were indeed deleted in the mutant strains (Fig. 6A-B). Our mutation of *tctD* appears to have potentially lowered the expression of *tctE*, albeit quite modestly (Fig. 6B). The Δ*tctE* strain showed normal or modestly elevated expression of *tctD* (Fig. 6A).

**Figure 6:**
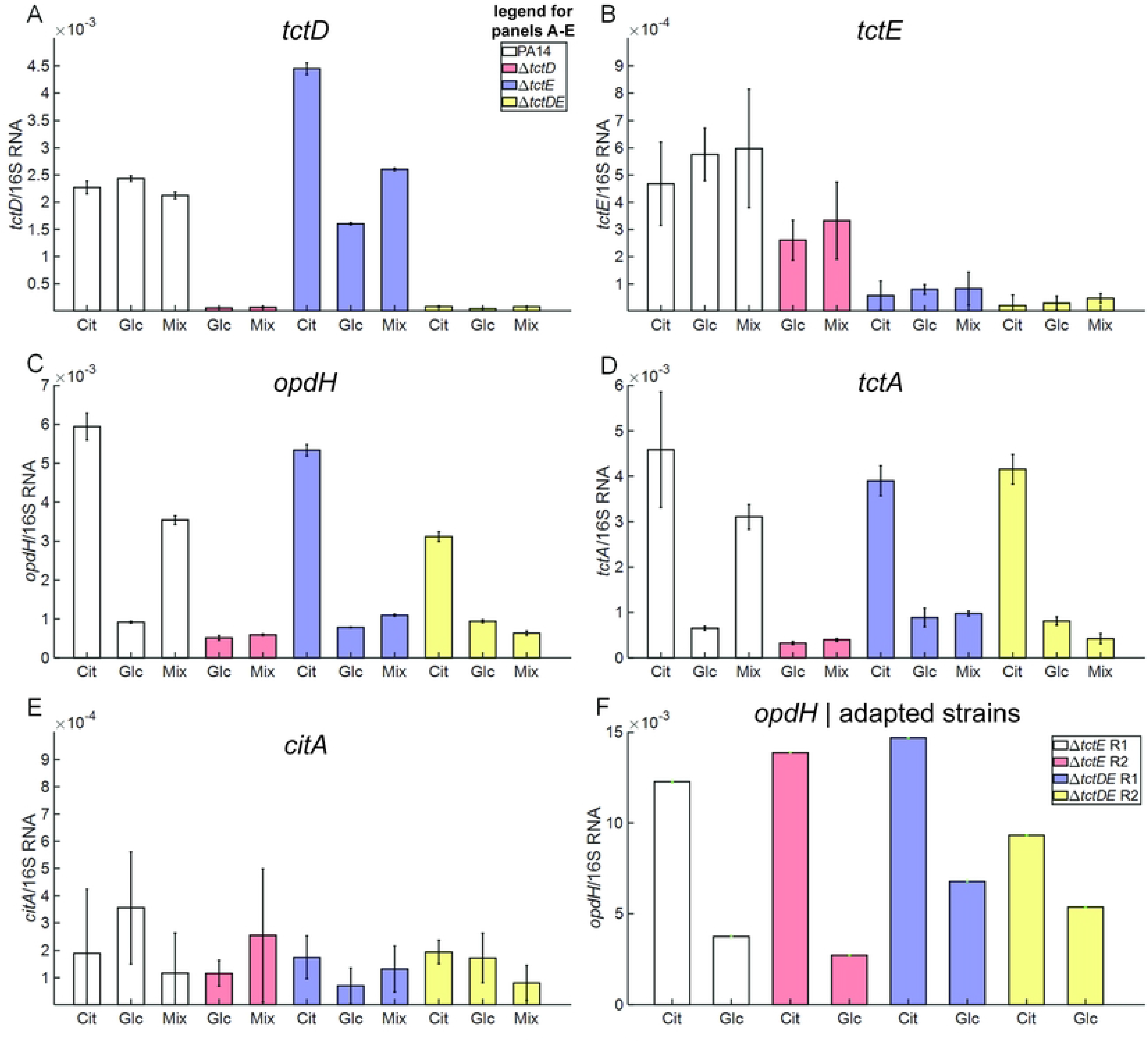
*tctDE* system mutants do not express *opdH* operon genes highly in the presence of citrate. RT-qPCR data showing gene expression per 16S rRNA unit. In panels (A)-(E), colors represent PA14 (white bars), Δ*tctD* (red bars), Δ*tctE* (blue bars), or Δ*tctDE* (yellow bars). In these panels, experiments were performed in citrate (cit), glucose (glc), or a mixture of citrate and glucose (mix). (A), *tctD* transcripts/16S rRNA; (B), *tctE* transcripts/16S rRNA; (C), *opdH* transcripts/16S rRNA; (D), *tctA* transcripts/16S rRNA; (E), *citA* transcripts/16S rRNA. (F) Expression of *opdH* in citrate-passaged mutants of Δ*tctE* (replicate 1, white bars; replicate 2, red bars) or Δ*tctDE* (replicate 1, blue bars; replicate 2, yellow bars).

We next assessed *opdH* expression, as our transcriptomic analysis identified it as a citrate-upregulated gene. Indeed, in PA14, expression of *opdH* is specifically induced by citrate, even in combination with glucose, as citrate is preferred over glucose by *P. aeruginosa* [1] (Fig. 6C). The Δ*tctD* mutant, however, failed to induce *opdH* in a citrate/glucose mixture, as expected given the failure of Δ*tctD* to grow in citrate (Fig. 5A). Similarly, *opdH* was not induced by citrate/glucose mixtures in the Δ*tctE* and Δ*tctDE* backgrounds (Fig. 6C). However, when grown in citrate alone, the Δ*tctE* and Δ*tctDE* strains regained expression of *opdH*, consistent with their (presumably adapted) growth (Fig. 6C). The same patterns of induction for the citrate/glucose mixture and for citrate alone were observed for *tctA* expression (Fig. 6D), suggesting that the whole *opdH-tctCBA* operon is expressed in *tct* mutants that gain the ability to grow on citrate.

Next, we tested whether citrate or the *tctDE* system are able to affect expression of *citA*. We expected *citA* to be affected by both, as control of *citA* by citrate and TctDE might explain how *P. aeruginosa* can encode a functional CitA standalone citrate transporter but *tctD* mutants cannot grow on citrate. Surprisingly, however, we found that *citA* was not induced on citrate (Fig. 6E). Moreover, *citA* was unaffected by the presence of the TcdDE TCS, and indeed appears to be barely transcribed at all on citrate or glucose (Fig. 6E), explaining why *citA* deletion has no growth phenotype on citrate (Fig. 1A).

Finally, we sought to formally test whether our citrate-adapted and LB-passaged Δ*tctE* and Δ*tctDE* strains had regained citrate-controlled *opdH* expression or whether they now constitutively expressed the *opdH-tctCBA* operon. Hence, we examined transcription of *opdH* in two passaged strains of each parental background grown in either citrate or glucose. We clearly saw that *opdH* was expressed less in glucose than in citrate across all samples, though the transcript per 16S rRNA count was 5 times higher than in the parental strain in glucose (Fig. 6C). In fact, even in glucose the passaged strains showed *opdH* expression levels that were roughly equal to the levels in citrate-grown wild-type cells (Fig. 6F vs. 6C). Collectively, these results suggest that the restoration of citrate growth in in the adapted Δ*tctDE* and Δ*tctE* strains is a product of much stronger basal *opdH-tctCBA* operon expression that nonetheless retains a degree of inducibility by citrate.

In an attempt to identify a genetic change conferring restored *opdH* operon expression and citrate growth, we sequenced the genomes of two independently passaged Δ*tctDE* strains and one Δ*tctE* strain and compared them to their respective parents. In all three passaged strains, SNPs not present in the parent were located in a region between a 16S rRNA gene and the annotated *sbcD* exonuclease gene. For both Δ*tctDE* passaged mutants, nucleotide 4957604 was changed from a T to C; in the Δ*tctE* passaged mutant there were two changes, 4957532 A to G and 4957549-50 GC to AT. In both of the Δ*tctDE* passaged mutants, there was additionally a frame shift in similar, C-terminal regions of the *pilY1* fimbrial biogenesis gene. As neither of these positions has any salient connection to citrate utilization, the significance of these SNPs with respect to the ability of the passaged strains to use tricarboxylates as carbon sources presently remains unclear.

### A Δ*tctD* mutant can grow in citrate if the *opdH* operon is expressed

We had observed that the Δ*tctD* and Δ*tctDE* mutants showed substantially reduced *opdH* transcript levels in medium containing both glucose and citrate and that the adapted strains showed elevated *opdH* expression. Thus, we hypothesized that loss of *tctD* blocks citrate utilization at least in part by abrogating expression of the *opdH-tctCBA-PA14_54580* operon, further implying that ectopic expression of one or more genes in the *opdH* operon might compensate for deletion of *tctD*. To test this notion, we inserted different genes from operon in pJN105, a replicative plasmid containing an arabinose-inducible multiple cloning site, to build a series of plasmids. We then transformed these plasmids into a Δ*tctD* strain and tracked the growth of the resulting strains in M9 containing citrate or *cis*-aconitate as the sole carbon source with and without the addition of 100 µM arabinose. We immediately observed that induction of the plasmid was not necessary to allow sufficient expression for growth on M9 citrate with at least some of the constructs (Fig. 7A). Importantly, a Δ*tctD* strain with empty pJN105 did not grow in citrate or *cis*-aconitate with or without arabinose but did grow in M9 with 3 mM glucose and gentamycin, indicating that the contents of the plasmid are important and demonstrating that the arabinose inducer does not serve as a growth-supporting carbon source (Supp. Fig. S5). All strains expressing *tctCBA* grew, and expression of additional genes in the operon improved growth relative to *tctCBA* alone (Fig. 7A-B). All the constructs except the full operon, which grew well even without induction, grew significantly better with arabinose induction than without (Fig. 7A-B), suggesting that that when the full operon is expressed, its expression level is not rate limiting for growth. We also found that, unlike in *E. coli*, expression of *citA* alone in Δ*tctD* does not enable growth in citrate (Fig. 7A-D), indicating that CitA alone cannot facilitate citrate transport in *P. aeruginosa*. We observed indistinguishable results in *cis*-aconitate (Fig. 7C and D, without and with arabinose, respectively). Collectively, our results argue that the citrate utilization defect in a Δ*tctD* mutant is caused by lack of *opdH* operon expression and that the core *tctCBA* genes are sufficient to restore citrate and *cis*-aconitate transport but are dramatically abetted by *opdH* and *54580* expression.

**Figure 7:**
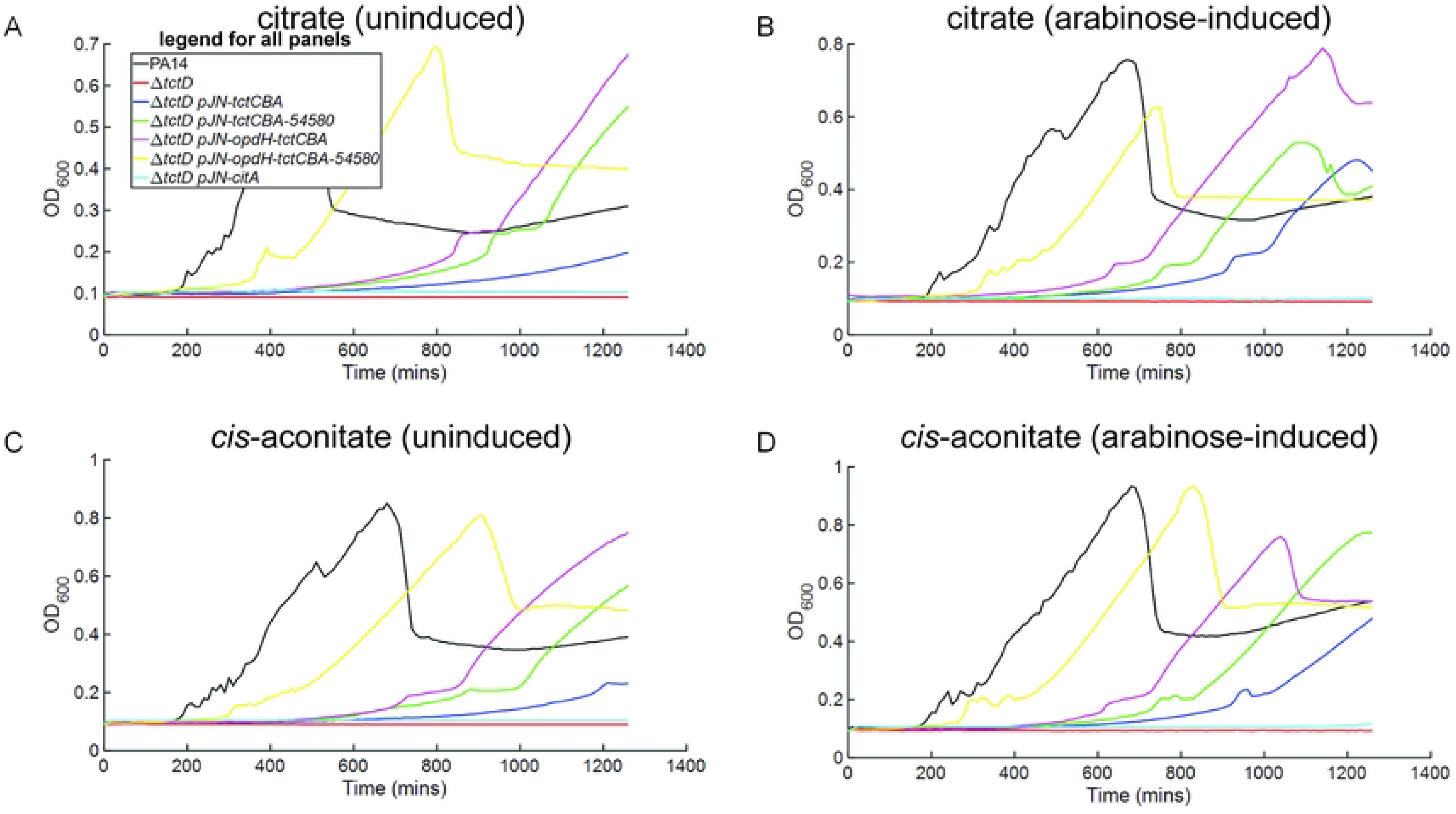
Ectopic expression of the *opdH* operon allows growth of Δ*tctD* on citrate and *cis*-aconitate. Growth curves showing OD_600_ vs. time for the Δ*tctD* mutant expressing portions of the *opdH* operon from an arabinose-inducible multiple cloning site on plasmid pJN105. Curves correspond to PA14 wild type (black curves); Δ*tctD* (red curves); Δ*tctD* pJN105-*tctCBA* (blue curves); Δ*tctD* pJN105-*tctCBA-PA14_54580* (green curves); Δ*tctD* pJN105-*opdH-tctCBA* (magenta curves); Δ*tctD* pJN105-*opdH-tctCBA-PA14_54580* (yellow curves); or Δ*tctD* pJN105-*citA* (cyan curves). Panels correspond to growth in M9 supplemented with 7.5 mM citrate as the sole carbon source without arabinose (A) or with 100 µM arabinose (B) or in 7.5 mM *cis*-aconitate as the sole carbon source without arabinose (C) or with 100 µM arabinose (D). Data are representative curves of at least 3 independent experiments.

## Discussion

Despite TCA cycle intermediates being preferred carbon sources for *P. aeruginosa* [8–10], the process of moving citrate molecules from the extracellular space to the cytoplasm has not been well understood in this organism. Previously, it was shown by Tamber *et al.* that transposon insertion in the *opdH* outer membrane porin-encoding gene results in a lack of growth on *cis*-aconitate [3]. However, further work by Tamber *et al.* failed to demonstrate transport of citrate or *cis*-aconitate by OpdH in a planar lipid bilayer experiment [11], suggesting that the transport mechanism does not rely on a single channel protein but may involve cooperation between outer-membrane and inner-membrane components. It has been thought that the *opdH* operon is repressed by TctD, the response regulator of the TctDE TCS [11, 14]. However, given the recently noticed growth defects of *tctDE* mutants on citric acid [15], if OpdH is the major transporter of citrate across the cell membrane, TctD would likely not be acting as a repressor, leading to questions as to how TctDE regulates citrate transport.

We first queried the importance of the OpdH porin in citrate and *cis*-aconitate transport by deleting it using a markerless method, in contrast to the Tamber *et al.* insertional method [11] that might have impacted downstream genes in the operon. We found (Fig. 1) that markerless deletion of *opdH* had no effect on growth in either carbon source. This agrees with the previous study [11] that OpdH is not crucial to transport. Combined with our data in Fig. 3A that *tctA* mutants lag in growth, we may reasonably infer that the earlier Tamber *et al.* study [3] may have inadvertently disrupted expression of the *tctCBA* section of the *opdH* operon in addition to the *opdH* gene, resulting in a strong growth defect on citrate. We confirm the notion that OpdH is not a standalone transport unit for citrate or *cis*-aconitate in Fig. 2, where expression of the *opdH* gene in *E. coli* fails to allow growth on either carbon source.

Having found an ortholog of the previously identified *Salmonella citA* gene, we next examined whether CitA is a major transporter of citrate. Deletion of *citA* had the same results as *opdH* deletion; however, deletion of both resulted in a growth lag on *cis*-aconitate but not citrate (Fig. 1). Expression of *citA* in *E. coli* MG1655 permitted some growth on citrate but substantially better growth on *cis-*aconitate (Fig. 2). The evidence from deletion in *P. aeruginosa* and complementation in *E. coli* suggests that perhaps CitA is primarily a transporter of *cis*-aconitate and has an additional weak ability to transport citrate. In RT-qPCR experiments we failed to observe significant transcription of *citA* in glucose, citrate, or a mix of both (Fig. 6E). This finding agrees with the idea that CitA is primarily a *cis*-aconitate transporter, and it will be interesting in future work to learn whether it is inducible by *cis*-aconitate.

As deletion of both *opdH* and *citA* did not yield a growth defect in citrate, we turned to transcriptomic comparisons to identify novel transporters. SCFM2 medium was used to simulate the infection environment of the cystic fibrosis lung, and because citrate should be preferred by *P. aeruginosa* before other carbon sources [8–10], we expected it to return a list of induced transporters despite the presence of other carbon sources. Encouragingly, we identified *opdH* and its downstream operon member *tctA*, which are known to be citrate-induced [11], in our list of upregulated genes. We selected *tctA,* as it is scantly studied and constitutes the predicted major transmembrane section of the TctCBA transporter, for markerless deletion and phenotypic screening. We also selected for deletion *rarD, PA14_72170*, and *PA14_18320*, which potentially encode transport systems. PA14_18320 and RarD are annotated as EamA family transporters, a group of proteins in the membrane drug/metabolite superfamily [23, 24]. PA14_72170 is marked as a CitMHS family transporter (Uniprot has it annotated as *citM*), a type of porin that transports divalent metal cations complexed with citrate [25].

In agreement with our hypothesis that these proteins are transporters for citrate, we found that a triple deletion of *tctA, 72170*, and either *rarD* or *18320* did not grow on citrate, suggesting that RarD and 18320 may work together. In addition, a Δ*tctA* single mutant lagged in growth on citrate, and additional deletion of *PA14_72170* exacerbated this lag, both in PA14 (Fig. 3) and Δ*citA* Δ*opdH* (Supp. Fig. S1) backgrounds. To our knowledge, ours is the first study to use a non-insertional, markerless deletion strategy to generate a *P. aeruginosa* strain that does not grow on citrate due to transporter knockout. However, the above results are insufficient to conclude that the predicted porins are themselves transporters of citrate and not another class of protein, such as membrane-bound regulators. To more specifically test their transport function, we expressed them from pTrc99A in *E. coli* MG1655. Among *tctA, 72170*, *rarD*, and *18320*, only *72170* transported citrate when individually expressed in *E coli* (Fig. 4 A, B). A ready explanation for why TctA alone failed to allow *E. coli* to grow on citrate is that it likely needs TctB and TctC to function; similarly, RarD and PA14_18320 may need one another or other partner proteins that are not present in *E. coli* to achieve transport.

How do these putative citrate transporters relate to transport of *cis-*aconitate? Deletion of *tctA* appears to completely abolish growth on aconitate (Fig. 3B, D), though not in a Δ*opdH* Δ*citA* background (Supp. Fig. S1). Strangely, deleting *PA14_72170* or *PA14_18320* in the Δ*tctA* strain restored growth on this carbon source, though the resulting mutant lags compared to the wild type. A triple Δ*tctA* Δ*72170* Δ*rarD* mutant, however, has almost no ability to grow on *cis*-aconitate (Fig. 3D). We are unsure how to interpret these results without further study but speculate that loss of TctA while keeping an intact OpdH could cause toxic periplasmic buildup of *cis*-aconitate if OpdH is the outer membrane porin and TctCBA the inner. The cellular locations of these porin systems are currently unknown but represent an important subject for future investigation. None of the newly identified transporters was able to allow growth of *E. coli* MG1655 on *cis*-aconitate (Fig. 4 C-D), implying that PA14_72170 is specific for citrate. We were also surprised to find that *PA14_54580*, the last member of the *opdH* operon, was inhibitory to *E. coli* growth when expressed in conjunction with the rest of the operon (Fig. 2C-D), but improved growth of a *P. aeruginosa* Δ*tctD* strain when expressed with the rest of the *opdH* operon. The 54580 protein was previously found to be 70% similar to AmoA of *Pseudomonas putida* [11], a membrane bound ammonia dioxygenase. It was suggested that because such enzymes can have fairly wide substrate specificity, 54580 could have a role in the metabolism of compounds transported by OpdH/TctCBA. If true, however, the data require that this function abet growth in *P. aeruginosa* but inhibit growth in *E. coli*; one possible mechanism for this difference is that *E. coli* does not possess a downstream metabolic pathway for a product of 54580 activity.

Having discovered putative new transporters, we next tried to understand the regulation of citrate transporter expression. It is known that Δ*tctDE* strains have a pH-independent growth defect when citric acid is added to M63-arginine medium [15] and that *P. aeruginosa* has elevated *opdH* expression in response to tricarboxylates [11], as confirmed by our transcriptomic analyses. We attempted to grow both a Δ*tctDE* strain and individual Δ*tctD* and Δ*tctE* mutants on citrate as the sole carbon source to understand the relative roles of these genes. We found that Δ*tctDE* and Δ*tctE* lag for about 2,000 and 1,400 minutes, respectively, while Δ*tctD* does not grow at all. This result does not fit the previous proposition that TctD is acting as a repressor of expression unless its loss causes such overexpression of transporters that the mutation is lethal. This unlikely scenario also conflicts with our data showing that the *tct* mutants grow normally on glucose or glucose/citrate mixes (Supp. Fig. S3) and our glucose/citrate mix qPCR data, which shows low transporter expression in the Δ*tctD* strain (Fig. 6C-D). Hence, the available evidence supports a model in which TctD is a positive regulator, not a repressor, of the *opdH* operon.

Because tandem loss of *tctE* and *tctD* does allow eventual growth on citrate, it is possible that loss of previously documented interactions between TctD and the PhoB response regulator [14] causes a TctE-dependent toxic effect. This would make sense, as TctD was shown to repress genes that are activated by PhoB. If a toxic gene were also tricarboxylate-inducible and the interaction TctE-dependent, such a strain would not grow on citrate or *cis*-aconitate. The Δ*tctD* strain was able to grow in citrate or *cis*-aconitate when elements of the *opdH* operon were expressed from pJN105 (Fig. 7), suggesting that if there is such a toxic effect then it can be remedied by allowing citrate transport into the cell. We speculate that without TctD there is a TctE-dependent way for citrate to enter the periplasm but not cross into the cytoplasm, resulting in a buildup that compromises cell growth. We also found that every element of the *opdH* operon contributes to improving the growth of this strain, indicating that while TctCBA may form the core transport machinery (its expression alone is sufficient to enable some growth), OpdH and PA14_54580 have supporting roles to play in transport or use of tricarboxylates. The fact that CitA functions alone as a transporter in *E. coli* but does not complement loss of *tctD* suggests that CitA requires a partner protein in *P. aeruginosa* that Δ*tctD* does not express – perhaps OpdH.

Our observation that the Δ*tctDE* and Δ*tctE* strains evolved heritable growth on citrate and regained the ability to express the *opdH* operon suggests that the use of citrate is highly important to *P. aeruginosa*, in agreement with its preference for this carbon source over saccharides. While the adapted strains exhibit some control of *opdH* expression by carbon source (Fig. 6F), there is also significantly higher transcription in cells grown in glucose compared to the parental strain or PA14 (Fig. 6F vs. Fig. 6C). These data suggest that while the adapted strains recover some level of *opdH* induction by the presence of citrate, the overall expression of the *opdH* operon is not as tightly regulated as in wild-type *P. aeruginosa*. Taking our results together, we argue that the TctDE TCS likely positively regulates citrate transport in *P. aeruginosa* rather than TctD acting as a repressor and that loss of TctDE can be compensated for via heritable mutations. Our efforts to find such mutations using whole genome sequencing have so far failed to yield any obvious candidates.

In summary, we have probed the transport of citrate and *cis*-aconitate in *P. aeruginosa* and identified proteins that are of interest in this process. We have clarified that the entire *opdH-tctCBA-PA14_54580* operon is impactful for tricarboxylate transport, with *tctCBA* as the key cross-membrane component. We have also shown that the TctDE TCS is important, but not strictly required, for growth on citrate and *cis*-aconitate and have supplied evidence for a possible mechanism of its control over transporter expression. Finally, we have shown that growth of *tctDE* mutants on citrate induces a heritable change of unknown character that enables high, possibly constitutive, expression of the *opdH* transport operon.

## Acknowledgments

The authors gratefully acknowledge funding from the NIGMS under grant number 1R35GM138018-01 and from the Oklahoma Center for Microbial Pathogenesis & Immunity (NIH CoBRE 1P20GM134973-01) to M.T.C. We would also like to thank Dr. Dave Dyer at the University of Oklahoma Health Sciences Center for his assistance in organizing the RNA sequencing experiments, and members of the Cabeen lab at Oklahoma State University for their helpful discussions and sharing of ideas. Finally, we acknowledge Dr. Zach Blount and Dr. Richard Lenski of Michigan State University for their advice and provision of strains; Dr. Tyrrell Conway at Oklahoma State University for his helpful discussion of metabolite transport and provision of *E. coli* strain MG1655; Dr. Boo Shan Tseng at the University of Nevada, Las Vegas, for providing plasmid pJN105; and Dr. Randy Morgenstein at Oklahoma State University for his provision of plasmid pTrc99A.

## Material and Methods

### Strains and growth conditions

*P. aeruginosa* and *E. coli* were grown in LB (Lennox) medium (10 g/L tryptone, 5 g/L yeast extract, 5 g/L NaCl) for overnight cultures. All liquid cultures were grown with shaking in 14 mL round-bottomed tubes at 180 RPM and 37°C unless otherwise specified. M9 medium was made according to the Cold Spring Harbor protocol [26] without the glucose, which was replaced with indicated carbon sources. In all cases, before experiments, cells were pelleted and washed twice in phosphate-buffered saline (PBS) pH 7.4 to remove any residual LB from the overnight culture. Cells were then diluted as indicated into the new growth medium used for the experiment. *E. coli* strains bearing pEXG2 or JN105 constructs were grown in an additional 20 µg/mL gentamycin; *P. aeruginosa* strains bearing pJN105 constructs were grown in 60 µg/mL gentamycin. *E. coli* bearing pTrc99A constructs were grown in 50 µg/mL carbenicillin, whereas *E. coli* bearing pCitT was grown in 50 µg/mL kanaymcin. Unless otherwise indicated, citrate and *cis*-aconitate were added to M9 at a concentration of 7.5 mM; glucose was added at a concentration of 3 mM. Mixtures of citrate and glucose were 7.5 mM citrate plus 3 mM glucose.

**Table 1:** *P. aeruginosa* strains used in this study. Parental lineages and details of plasmids and *E. coli* cloning/conjugation strains are given in Supporting Information. Ge, gentamycin.

| Strain Number | Genotype | Characteristics | Source or reference |
| --- | --- | --- | --- |
| MTC1 | PA14 | Laboratory wild type of *P. aeruginosa* PA14 | [27] |
| MTC2550 | $\Delta citA$ | PA14 with markerless *citA* deletion | This study |
| MTC2551 | $\Delta opdH$ | PA14 with markerless *opdH* deletion | This study |
| MTC2552 | $\Delta citA \Delta opdH$ | PA14 with markerless *opdH* and *citA* deletions | This study |
| MTC2553 | $\Delta tctA$ | PA14 with markerless *tctA* deletion | This study |
| MTC2554 | $\Delta rarD$ | PA14 with markerless *rarD* deletion | This study |
| MTC2555 | $\Delta PA14\_18320$ | PA14 with markerless *PA14_18320* deletion | This study |
| MTC2556 | $\Delta PA14\_72170$ | PA14 with markerless *PA14_72170* deletion | This study |
| MTC2557 | $\Delta tctD$ | PA14 with markerless *tctD* deletion | This study |
| MTC2558 | $\Delta tctE$ | PA14 with markerless *tctE* deletion | This study |
| MTC2559 | $\Delta tctDE$ | PA14 with markerless *tctDE* deletion | This study |
| MTC2560 | $\Delta tctE$ passaged | $\Delta tctE$ passaged in M9-citrate, streaked on LB, and then frozen from LB overnight culture. | This study |
| MTC2561 | $\Delta tctDE$ passaged R1 | $\Delta tctDE$ passaged in M9-citrate, streaked on LB, and then frozen from LB overnight culture.<br>Replicate 1. | This study |
| MTC2562 | $\Delta tctDE$ passaged R2 | $\Delta tctDE$ passaged in M9-citrate, streaked on LB, and then frozen from LB overnight culture.<br>Replicate 2. | This study |
| MTC2563 | $\Delta tctD$ pJN105 empty vector | PA14 $\Delta tctD$ carrying pJN105 empty vector, Ge<sup>R</sup> | This study |
| MTC2564 | $\Delta tctD$ pJN105-*tctCBA* | PA14 $\Delta tctD$ carrying pJN105 encoding the *tctCBA* gene cluster under arabinose inducible promoter, Ge<sup>R</sup> | This study |
| MTC2565 | $\Delta tctD$ pJN105-<br>*opdH-tctCBA* | PA14 $\Delta tctD$ carrying<br>pJN105 encoding the<br>*opdH-tctCBA* gene cluster<br>under arabinose inducible<br>promoter, Ge<sup>R</sup> | This study |
| MTC2566 | $\Delta tctD$ pJN105-<br>*tctCBA-*<br>*PA14_54580* | PA14 $\Delta tctD$ carrying<br>pJN105 encoding the<br>*tctCBA-PA14_54580* gene<br>cluster under arabinose<br>inducible promoter, Ge<sup>R</sup> | This study |
| MTC2567 | $\Delta tctD$ pJN105-<br>*opdH-tctCBA-*<br>*PA14_54580* | PA14 $\Delta tctD$ carrying<br>pJN105 encoding the<br>entire *opdH-tctCBA-*<br>*PA14_54580* operon<br>under arabinose inducible<br>promoter, Ge<sup>R</sup> | This study |

**Table 2:** *E. coli* strains used in experiments in this study. Cb, carbenicillin; Ge, gentamycin; Km, kanamycin.

| Strain Number | Genotype | Characteristics | Source or reference |
| --- | --- | --- | --- |
| MTC2568 | MG1655 | Wild type K-12<br>derivative *E. coli* | Yale Genetic Stock<br>Culture Collection<br>CGSC#6300 |
| MTC2569 | MG1655 pCitT | MG1655 bearing *E. coli* CitT citrate<br>transporter, Km<sup>R</sup> | This study |
| MTC2570 | MG1655 pTrc99A-<br>*citA* | MG1655 bearing<br>pTrc99A plasmid<br>with *citA* inserted at<br>*lac* controlled<br>multiple cloning<br>site, Cb<sup>R</sup> | This study |
| MTC2571 | MG1655 pTrc99A-<br>*opdH* | MG1655 bearing<br>pTrc99A plasmid<br>with *opdH* inserted<br>at *lac* controlled<br>multiple cloning<br>site, Cb<sup>R</sup> | This study |
| MTC2572 | MG1655 pTrc99A-<br>*tctCBA* | MG1655 bearing<br>pTrc99A plasmid<br>with *tctCBA*<br>inserted at *lac* controlled multiple cloning site, Cb<sup>R</sup> | This study |
| MTC2573 | MG1655 pTrc99A-<br>*opdH-tctCBA* | MG1655 bearing pTrc99A plasmid with *opdH-tctCBA* inserted at *lac* controlled multiple cloning site, Cb<sup>R</sup> | This study |
| MTC2574 | MG1655 pTrc99A-<br>*tctCBA-PA14_54580* | MG1655 bearing pTrc99A plasmid with *tctCBA-PA14_54580* inserted at *lac* controlled multiple cloning site, Cb<sup>R</sup> | This study |
| MTC2575 | MG1655 pTrc99A-<br>*opdH-tctCBA-PA14_54580* | MG1655 bearing pTrc99A plasmid with *opdH-tctCBA-PA14_54580* inserted at *lac* controlled multiple cloning site, Cb<sup>R</sup> | This study |
| MTC2576 | MG1655 pTrc99A-<br>*tctA* | MG1655 bearing<br>pTrc99A plasmid<br>with *tctA* inserted at<br>*lac* controlled<br>multiple cloning<br>site, Cb<sup>R</sup> | This study |
| MTC2577 | MG1655 pTrc99A-<br>*PA14_72170* | MG1655 bearing<br>pTrc99A plasmid<br>with *PA14_72170*<br>inserted at *lac*<br>controlled multiple<br>cloning site, Cb<sup>R</sup> | This study |
| MTC2578 | MG1655 pTrc99A-<br>*PA14_18320* | MG1655 bearing<br>pTrc99A plasmid<br>with *PA14_18320*<br>inserted at *lac*<br>controlled multiple<br>cloning site, Cb<sup>R</sup> | This study |
| MTC2579 | MG1655 pTrc99A-<br>*rarD* | MG1655 bearing<br>pTrc99A plasmid<br>with *rarD* inserted<br>at *lac* controlled multiple cloning<br>site, Cb<sup>R</sup> | This study |

### Strain construction

*E. coli* strains bearing plasmids were generated by standard chemical competence transformation with Gibson-assembled plasmids (see Supporting Information). *P. aeruginosa* allelic replacement mutants were generated using plasmid pEXG2 [28] containing flanking homologous regions of the gene to be deleted, which were amplified using standard PCR. Plasmids were inserted into *P. aeruginosa* PA14 by conjugation with *E. coli* SM10 on LB agar (see Supporting Information).

### Growth curve experiments

For the automated measurement of growth curves, cells were washed in PBS from overnight culture as described above. M9 was prepared with either 3 mM glucose, 7.5 mM citrate, or 7.5 mM *cis*-aconitate as indicated. Cells were diluted 100-fold into a clear 96 well polystyrene plate (Corning, Inc.) and covered with a lid. This was then incubated at 37°C in a BioTek Synergy H1 plate reader (BioTek, CA, USA) and the OD_600_ measured every ten minutes after a 2 second shake.

For the Δ*tct* mutant back-dilution experiment, cells were first grown in M9 with 7.5 mM citrate until the culture was turbid. 1 mL of the culture was then transferred to a 1.5 mL microcentrifuge tube and the cells washed twice in PBS by centrifugation and resuspension. The resulting cells were 100-fold back-diluted into shaking flasks containing M9 with 7.5 mM citrate. Growth was monitored by measurement of the OD_600_ using cuvettes in a spectrophotometer.

### RT-qPCR experiments

For RT-qPCR, cells in biological triplicate overnight cultures were diluted 50-fold from 2x PBS wash into 50 mL M9 with either 3 mM glucose, 7.5 mM citrate, or 3 mM glucose plus 7.5 mM citrate. Cells were allowed to grow to an OD of between 0.198-0.397 (mid-exponential phase growth) before centrifugation. The pellet was then re-suspended in 250 µL of TE buffer (10 mM Tris pH 8.0, 1 mM EDTA) supplemented with 10 mg/mL lysozyme from chicken egg white and allowed to incubate at room temperature for 15 mins. Following incubation, the RNA extraction protocol from New England Biolabs’ Monarch RNA Extraction kit (NEB, MA, USA) was followed. The purified RNA was quantified using UV spectrophotometry a,nd 500 ng RNA was reverse transcribed using the RevertAid RT Reverse Transcription Kit (ThermoFisher, MA, USA). The resulting cDNA was diluted 25-fold in water and 1 µL of this used in each 10 µL qPCR reaction. Reactions were carried out in BioRad semi-skirted qPCR plates covered with an adhesive film (BioRad, CA, USA) using the Promega GoTaq BRYT Green^®^ dye-based qPCR kit (Promega, WI, USA) with 200 nM concentration of the appropriate primers. The plate was centrifuged at 500 RPM in a Sorvall ST 40R centrifuge equipped with a TX-1000 rotor for 2 minutes. This was then processed in a CFX96 Touch real-time PCR cycler (BioRad, CA, USA). Initial denaturation was performed for 2 minutes at 95°C, and cycles were performed using a 15 second denaturation step at 95°C followed by 1 minute annealing and extension at 60°C.

Results of the qPCR were determined using the standard curve method. Standards were produced by gel-purifying an ordinary PCR made using the qPCR primers and PA14 genomic DNA. The result was quantified spectrophotometrically and the average weight per base of the strand used to dilute it to a concentration of 10^9^ copies/µL. This was then serially 10-fold diluted to generate a standard curve down to 10^2^ copies/µL. These standards were included in the qPCR plate and a plot of log(copy number) vs. C_q_ (the number of cycles needed to reach the fluorescence quantification threshold) used to determine the relationship between C_q_ and the copy number of DNA strands. The standard was then used to infer how many copies of a particular reverse-transcribed RNA were present in the cDNA samples. Each of these was then divided by the number of 16S rRNA copies to normalize to a housekeeping gene. Error was calculated by taking the standard error in the mean of two technical replicates for each biological replicate, both for the 16S measurement and the queried genes. These errors were then propagated forward into the quotient using standard error propagation methods.

### RNA sequencing

*RNA* for sequencing was extracted as for the RT-qPCR experiments. The RNA was then depleted of ribosomes using the Illumina RiboZero kit (Illumina, CA, USA). Depleted samples were submitted for Illumina sequencing at the University of Oklahoma Health Sciences Center core facility in Oklahoma City, OK, USA. Sequence mapping and analysis were performed at the Oklahoma University Health Sciences Center Laboratory for Molecular Biology and Cytometry Research using CLC software.

### Whole genome sequencing

Samples for whole genome sequencing were grown overnight in LB as above. 1 mL of cells was then processed using the Promega Wizard^®^ genomic DNA preparation kit (Promega, WI, USA) to obtain genomic DNA. The DNA was then shipped on dry ice to Novogene (Novogene Corporation Inc., CA, USA) for quality control, library preparation, and sequencing.

